# SARS-CoV-2 infects and induces cytotoxic effects in human cardiomyocytes

**DOI:** 10.1101/2020.06.01.127605

**Authors:** Denisa Bojkova, Julian U. G. Wagner, Mariana Shumliakivska, Galip S. Aslan, Umber Saleem, Arne Hansen, Guillermo Luxán, Stefan Günther, Minh Duc Pham, Jaya Krishnan, Patrick N. Harter, Utz Ermel, Achilleas Frangakis, Andreas M. Zeiher, Hendrik Milting, Jindrich Cinatl, Andreas Dendorfer, Thomas Eschenhagen, Sandra Ciesek, Stefanie Dimmeler

## Abstract

**Background:** The coronavirus disease 2019 (COVID-19) is caused by severe acute respiratory syndrome coronavirus 2 (SARS-CoV-2) and has emerged as global pandemic. SARS-CoV-2 infection can lead to elevated markers of cardiac injury associated with higher risk of mortality in COVID-19 patients. It is unclear whether cardiac injury may have been caused by direct infection of cardiomyocytes or is mainly secondary to lung injury and inflammation. Here we investigate whether human cardiomyocytes are permissive for SARS-CoV-2 infection.

**Methods:** Infection was induced by two strains of SARS-CoV-2 (FFM1 and FFM2) in human induced pluripotent stem cells-derived cardiomyocytes (hiPS-CM) and in two models of human cardiac tissue.

**Results:** We show that SARS-CoV-2 infects hiPS-CM as demonstrated by detection of intracellular double strand viral RNA and viral spike glycoprotein protein expression. Increasing concentrations of virus RNA are detected in supernatants of infected cardiomyocytes, which induced infections in CaCo-2 cell lines documenting productive infections. SARS-COV-2 infection induced cytotoxic and pro-apoptotic effects and abolished cardiomyocyte beating. RNA sequencing confirmed a transcriptional response to viral infection as demonstrated by the up-regulation of genes associated with pathways related to viral response and interferon signaling, apoptosis and reactive oxygen stress. SARS-CoV-2 infection and cardiotoxicity was confirmed in a iPS-derived human 3D cardiosphere tissue models. Importantly, viral spike protein and viral particles were detected in living human heart slices after infection with SARS-CoV-2.

**Conclusions:** The demonstration that cardiomyocytes are permissive for SARS-CoV-2 infection in vitro warrants the further in depth monitoring of cardiotoxic effects in COVID-19 patients.

**Clinical Perspective:** *What is New?:*

- This study demonstrates that human cardiac myocytes are permissive for SARS-CoV-2 infection.
- The study documents that SARS-CoV-2 undergoes a full replicatory circle and induces a cytotoxic response in cardiomyocytes.
- Infection was confirmed in two cardiac tissue models, including living human heart slices.

*What are the Clinical Implications?:*

- The study may provide a rational to explain part of the cardiotoxicity observed in COVID-19 patients
- The demonstration of direct cardiotoxicity induced by SARS-CoV-2 warrants an in depth further analysis of cardiac tissue of COVID-19 patients and a close monitoring for putative direct cardiomyocyte injury.
- The established models can be used to test novel therapeutic approaches targeting COVID-19.

## Introduction

The coronavirus disease 2019 (COVID-19) is caused by severe acute respiratory syndrome coronavirus 2 (SARS-CoV-2) and has emerged as global pandemic. SARS-CoV-2 mainly invades alveolar epithelial cells and causes adult respiratory distress syndromes. COVID-19 is associated with myocardial injury, as assessed by increased troponin T and NT-proBNP levels accompanying increased cardiovascular symptoms in a significant number of SARS-CoV-2 infected patients^1,2^. Elevated levels of cardiac injury markers were associated with higher risk of in-hospital mortality in COVID-19 patients^3^. In patients showing clinical deterioration during COVID-19, left ventricular systolic dysfunction was noted in approximately 20 % of patients according to a most recent study^4^. In addition, patients with underlying cardiovascular disease represent a significant proportion of patients, who may suffer from severe courses after COVID-19 infections^5^. However, it is unclear whether elevated biomarkers of cardiac injury and long term effects on the cardiovascular system are directly caused by viral infection of cardiac tissue or are secondary to hypoxia and systemic inflammation during complicated COVID-19 courses. Earlier studies with cardiac tissue samples did not find evidence for viral particles of the first SARS corona virus SARS-CoV^6^, but other studies suggest that the Middle East respiratory syndrome-related coronavirus (MERS-CoV), which has similar pathogenicity as SARS-CoV-2, can cause acute myocarditis and heart failure^7^. Moreover, substantial amounts of viral SARS-CoV-2 RNA was detected in human hearts of COVID-19 patients^8^.

Single cell RNA sequencing and histological analyses demonstrated that human cardiomyocytes express the putative SARS-CoV-2 receptor angiotensin converting enzyme 2 (ACE2), particularly in patients with cardiovascular diseases^9,10^ suggesting that cardiomyocytes could be targeted by SARS-CoV-2.

Therefore, we investigated whether SARS-CoV-2 infects human induced pluripotent stem cell-derived cardiomyocytes in culture and in two models of human cardiac tissue including human heart slices *in vitro*.

## Methods

### Cell Culture

hiPS-CM of two donors were obtained with an embryoid body-based protocol as described^11^. Cardiospheres were generated by adapting a previously described protocol^12^ using hiPS cells. Living human heart slices (300 μm) were generated and cultured as described^9^

### Viral infection

SARS-CoV-2-FFM1 and FFM2 were isolated and propagated in Caco-2 cells as described ^13,14^. The viral stock was diluted to desired MOI in medium containing 1% fetal bovine serum and incubated with cells for 2 h. Then the infectious inoculum was removed and cells were supplemented with the respective culture medium^11^. Cardiospheres were cultured with 25μl of viral stock (1.10^7^ TCID50/ml) and living human heart slices were incubated with 200μl of viral stock (1.10^7^ TCID50/ml) for three to five days.

Quantification of SARS-CoV RNA in cell culture supernatants was performed as previously described^14^. For detection of viral titer, hiPS-CM were infected for 2 h, the infection medium was replaced, and supernatants were collected 48h post infection and used to infect confluent layers of CaCo-2 cells in 96-well plates. Cytopathogenic effects were assessed visually 48 h after infection. The infectious titer was determined as TCID50/ml.

For further details, see Online Data Supplemental “Expanded Methods”.

## Results

### Expression of receptor and co-receptor

We first addressed if human induced pluripotent stem cell-derived cardiomyocytes (hiPS-CM) showed the expression of the SARS-CoV-2 receptors ACE2 and the serine proteases TMPRSS2 and cathepsins, which mediate priming of the viral S-protein^15^. ACE2 and cathepsins CTSB and CTSL were well expressed on mRNA level in hiPS-CM, whereas TMPRSS2 was detected only at very low levels by RNA sequencing (**Figure 1a**). Quantitative RT-PCR confirmed the expression of ACE2, but TMPRSS2 was below the detection level (**Figure 1b**). ACE2 protein expression was confirmed by immunostainings using two different antibodies (**Figure 1c**, **Online Supplement Figure I**) documenting that human cardiomyocytes possess the so far described receptor and activators necessary for effective SARS-CoV-2 infection.

**Figure 1:**
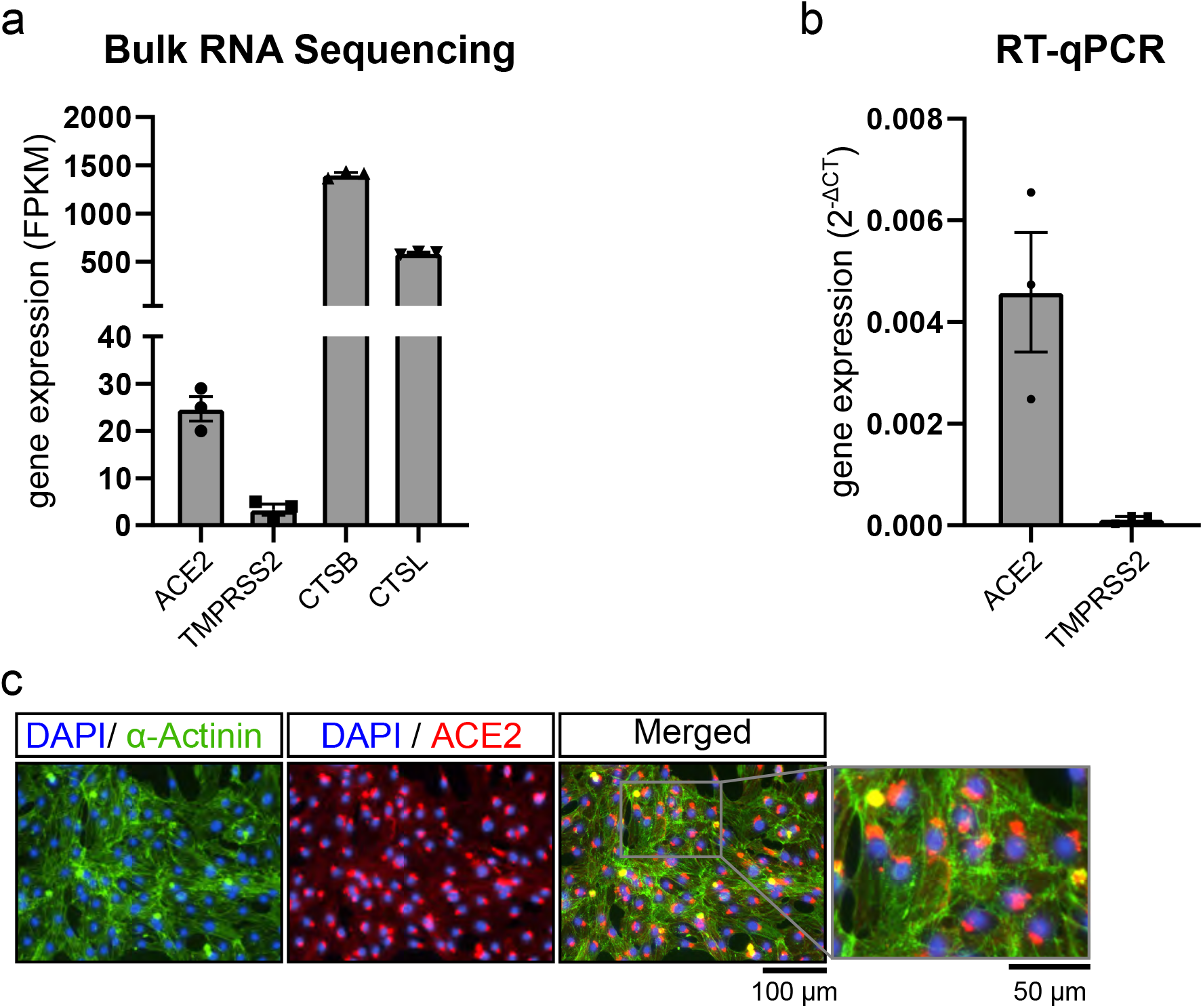
Expression of SARS-CoV-2 receptors and co-activators in hiPS-derived cardiomyocytes. **a**, Expression in RNA sequencing data of hiPS-CM (n=3). B, Confirmation by qRT-PCR (n=3), **c**, ACE2 protein expression was detected by antibody (Abcam) in hiPS-CMs. Cells were counterstained by α–sarcomeric actinin and DAPI. A representative experiment out of n>6 with two different iPS-CM donors is shown.

### hiPS-CM are infected by SARS-CoV-2

To test if hiPS-CM are directly targeted and are permissive for SARS-CoV-2 infection, hiPS-CM were infected with isolates of SARS-CoV-2^13^ (**Figure 2a**). SARS-CoV-2 infected hiPS-CM showed increased intracellular double stranded virus RNA as demonstrated by immunostaining (**Figure 2b**). Viral RNA in supernatants was further assessed by PCR and was dose- and time-dependently increased after infection with SARS-CoV-2 (**Figure 2c-e**). Consistently, the expression of the viral spike glycoprotein protein was detected in a time- and dose dependent manner after infection with different strains of the virus (FFM1 and FFM2^13^) (**Figure 2f-i**). Control experiments confirmed spike protein expression in α-sarcomeric actinin-expressing cardiomyocytes (**Figure 2i**). The supernatant of hiPC-CM contained fully infectious virus, as demonstrated by titration in CaCo-2 cells (**Figure 2j**) indicating that the virus undergoes full replicatory cycles in hiPS-CM. Functionally, SARS-CoV-2 infection reduced cell counts (**Figure 2k).** and augmented apoptosis in hiPS-CM (**Figure 2l**). Interestingly, the frequency of beating was significantly augmented at 24h to 48h post infection, but was abolished at later time points (**Figure 2m**). Profound cytopathogenic effects were visible at later time points (96 h) (**Figure 2h**). RNA sequencing demonstrated that infected hiPS-CM showed a strong transcriptional response to viral infection including interferon activation (**Figure 2n**) and signatures of apoptosis and oxidative stress (**Figure 2o**).

**Figure 2:**
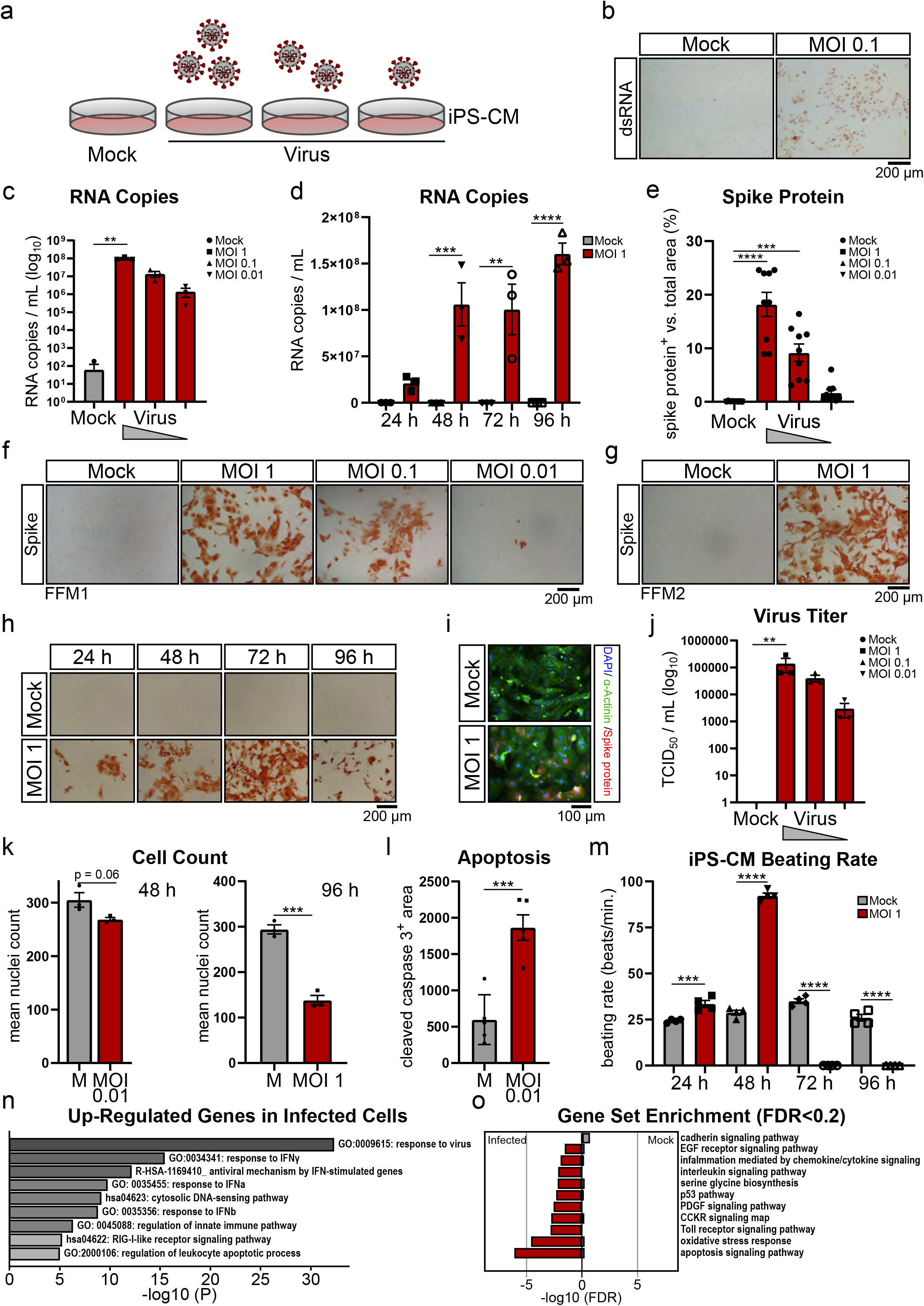
SARS-CoV-2 infects iPS-derived cardiomyocytes. **a**, Design of experiment. **b**, Immunostaining of double-stranded RNA (dsRNA) in hiPS-CMs after infection with SARS-CoV-2 FFM1 for 48 h. **c,d**, SARS-CoV-2 RNA was measured by qRT-PCR in the supernatant of infected hiPS-CM (d: 48 h, e: MOI 1). **e-h**, Spike glyocoprotein was measured by immunohistochemistry after infection with isolate SARS-CoV-2 FFM1 (f-g,i) or SARS-CoV-2 FFM2 (g). **i**, Counterstainings with α-sarcomeric actinin confirmed expression of spike protein in hiPS-CM (48h). **j,** Infectious virus in supernatants from infected hiPS-CM was determined by titration in Caco-2 cells 48 h post infection. **k,** Number of cells (DAPI+ nuclei) after 48h (MOI 0.01) or 96h (MOI 1) of infection (V) vs. mock (M) control. **l**, Quantification of cleaved caspase-3+ area after 48 h of infection (V) vs. mock (M). **m**, Beating rate of hiPS-CMs. **n,o**, RNA sequencing of mock or SARS-CoV-2 infected hiPC-CM (MOI 1, 48 h), n=3 each. **n**, GO term analysis (Metascape) of top up-regulated genes (>5 fold, FDR<0.2). **o**, Top10 enriched terms in PANTHER database for DEGs, FDR<0.2. Data are mean+SEM analyzed by using two-sided, unpaired t-test (k,l), ordinary one-way ANOVA with post hoc Tukey’s (e, m), Dunn’s (f) comparison or Kruskal-Wallis test with Dunn’s comparison (d, k). MOI: multiplicity of infection, **p<0.01, ***p<0.001, ****p<0.0001. All data are n≥3.

### SARS-CoV-2 infects cardiomyocytes in three dimensional cardiac tissue

Next, we determined if SARS-CoV-2 infects cardiomyocytes in a three dimensional tissue environment using human cardiospheres generated by hiPS-cells, which are generated by a modified previously published protocol^12^ (**Figure 3a-b**). SARS-CoV-2 time-dependently affected beating frequency of cardiospheres with a profound inhibition at 5 days post infection (**Figure 3c**). At 5 days post infection, cardiospheres showed a reduced size (**Figure 3d**) consistent with the induction of cell death. SARS-CoV-2 infection was further documented by spike protein staining (**Figure 3d**).

**Figure 3:**
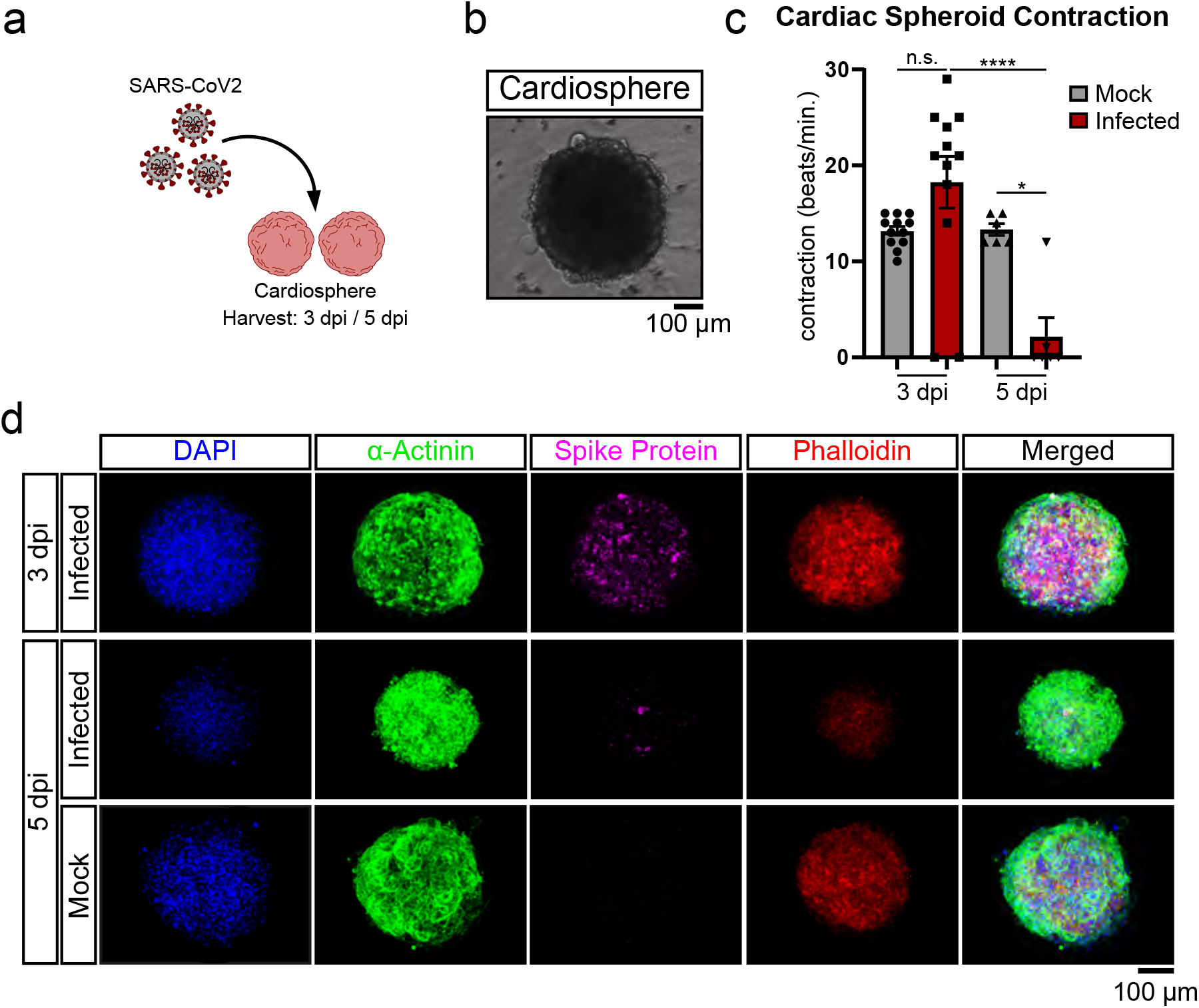
SARS-CoV-2 infects iPS-derived human cardiospheres. **a**, Study design of cardiosphere infection, **b**, hiPS-derived cardiosphere (light microscope image). **c**, Beating frequency of cardiospheres after 3 and 5 days post infection (dpi). **d,** Expression of spike glycoprotein at 3 and 5 days post infection (dpi). Data were statistically assessed using one-way ANOVA with post hoc Tukey’s test. n.s. = not significant, **p<0.05, ****p<0.0001.

Finally, we addressed whether SARS-CoV-2 infects human heart tissue by using living human cardiac tissue slices, which were obtained from explanted hearts^16^ (**Figure 4a-f**). Here, increased spike protein expression was shown in four different samples derived from three donors (**Figure 4c-e**). Infection was associated with morphological signs of tissue injury such as areas with loss of α-saromeric actinin signal and disorganized structure compared to homogenous mock controls (**Figure 4c**). Spike protein expression was detected in a-sacromeric actinin positive cardiomyocytes (**Figure 4d**). The virus was further identified in infected human heart tissue in cardiomyocytes by electron microscopy (**Figure 4f**). Of note, stages of the entire replicatory cycle were detected (**Figure 4f**, right panel)

**Figure 4:**
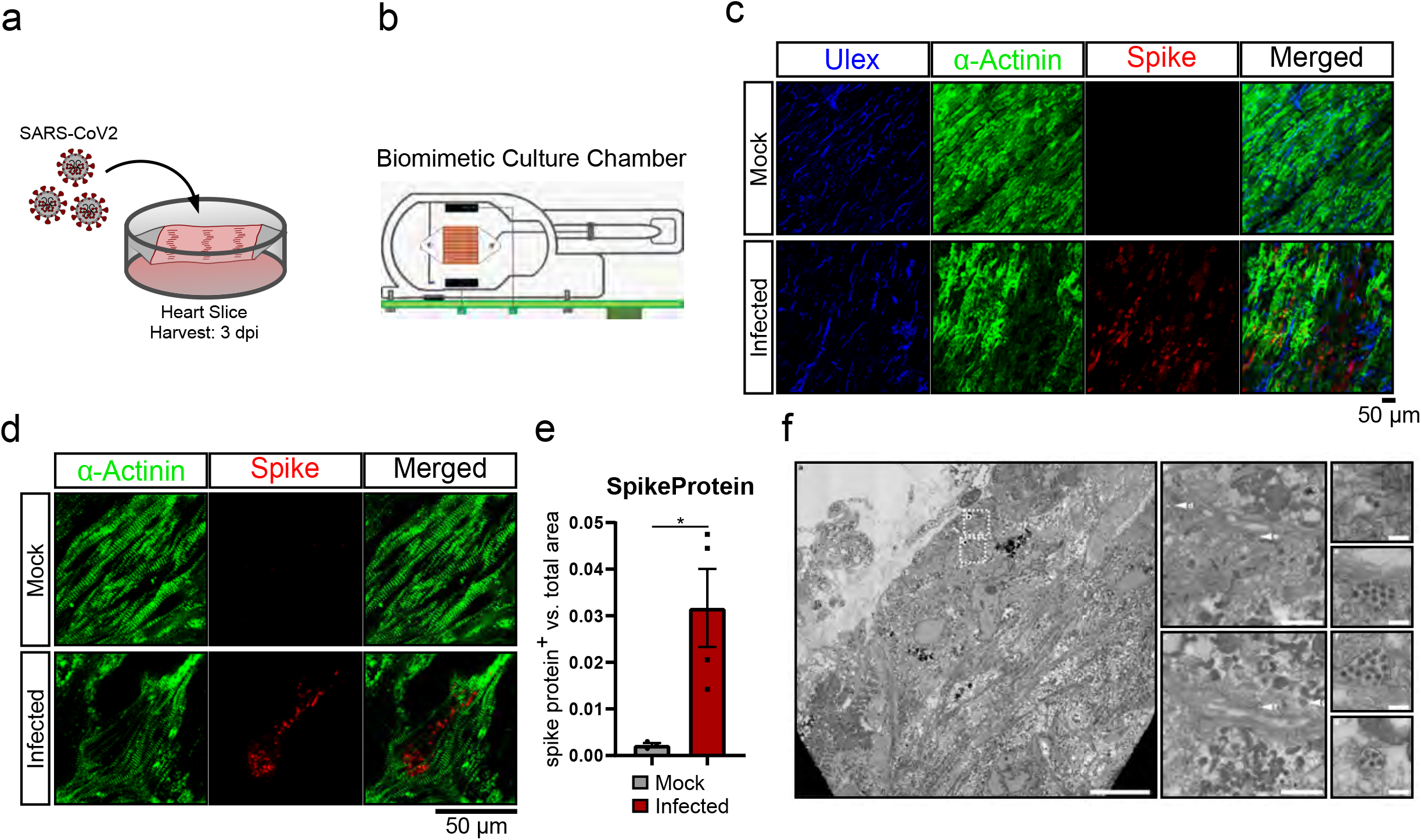
SARS-CoV-2 infects living human cardiac tissue slices. **a-c**, Infection of human cardiac tissue slices of three donors. **b**, Scheme of experiment. **c-e**, Spike protein expression in infected human heart slices (4 samples of three donors). Quantification of individual images of each donor is shown in **e**. Data were statistically assessed with unpaired T-test with Welch’s correction. *p<0.05. Representative images are shown in **c**, higher magnification of a representative cardiomyocyte expressing spike protein is shown in **d**. **f**, Electron microscopy of infected human cardiac tissue slices. Arrows indicate virus particles.

## Discussion

Together, SARS-CoV-2 can infect human cardiomyocytes in culture as well as in two different models of cardiac tissue. Infection was documented by various read outs including intracellular viral double strand RNA and spike protein expression, as well as extracellular viral RNA. The virus was further detected by electron microscopy in cells of the infected human heart slices. Importantly, functional virus could be isolated in supernatants of infected cardiomyocytes documenting that SARS-CoV-2 undergoes a full replicatory cycle. Of note, viral infection was confirmed with cells of two different hiPS-donors and two viral strains. Whereas the receptor ACE2 was well expressed on mRNA and protein level, the previously described protease activator TMPRSS2 was very lowly expressed in hiPS-CM, suggesting that activation of the S-protein may be mediated by other serine proteases such as cathepsins, which also can mediate viral activation^15^. Viral infection was associated with cytotoxic effects and inhibition of beating of cardiomyocytes in our in vitro cultures and cardiospheres suggesting a potential detrimental effect of SARS-CoV-2 infection on the human heart.

SARS-CoV-2 elicits a typical transcriptional response to viral infection including the activation of interferon pathways. Since we recently demonstrated that SARS-CoV-2 infection deregulates pathways involved in ER stress and protein homeostasis^14^, one may hypothesize that viral infection may induce ER stress leading to prolonged unfolded protein response and subsequent alteration in calcium homeostasis and cardiomyocyte cell death ^17^.

Although there is compelling evidence that patients suffering from COVID-19 show profound elevations of cardiac injury biomarkers and deteriorated right and left ventricular cardiac function, a direct viral infection of cardiomyocytes by SARS-CoV-2 has not been demonstrated so far. Viral RNA, however, has been detected at significant levels in cardiac tissue^8^ and cardiomyocyte infection may occur during conditions of vascular leakage and tissue inflammation. The lack of clear-cut detection of virus in cardiomyocytes so far, may be related the difficulties in findings the areas of infection and the rapid induction of cell death and clearance of the infected cells.

The profound effects of SARS-CoV-2 on human cardiomyocytes observed in the present in vitro study warrant the further in depth monitoring of direct cardiac effects in COVID-19 patients. The used 3D tissue models may serve as experimental model for testing the effects of coronavirus infection and biology in the heart and developing therapeutic strategies.

## Abbreviations

ACE2: angiotensin converting enzyme 2
COVID-19: coronavirus disease 2019
ER: endoplasmic reticulum
hiPS-CM: human induced pluripotent cardiomyocytes
SARS-CoV: severe acute respiratory syndrome coronavirus

## Acknowledgement

The authors thank Tatjana Starzetz (Neurological Institute (Edinger Institute), University Frankfurt) as well as Marion Basoglu (Biological Sciences, University Frankfurt) for EM Sample Preparation.

## Disclosures section

The authors have nothing to disclose

## Supplemental Materials

Expanded Methods

Online-only Figures I – IV

